# Syncytia formation by SARS-CoV-2 infected cells

**DOI:** 10.1101/2020.07.14.202028

**Authors:** Julian Buchrieser, Jeremy Dufloo, Mathieu Hubert, Blandine Monel, Delphine Planas, Maaran Michael Rajah, Cyril Planchais, Françoise Porrot, Florence Guivel-Benhassine, Sylvie Van der Werf, Nicoletta Casartelli, Hugo Mouquet, Timothée Bruel, Olivier Schwartz

## Abstract

Severe cases of COVID-19 are associated with extensive lung damage and the presence of infected multinucleated syncytial pneumocytes. The viral and cellular mechanisms regulating the formation of these syncytia are not well understood. Here, we show that SARS-CoV-2 infected cells express the viral Spike protein (S) at their surface and fuse with ACE2-positive neighbouring cells. Expression of S without any other viral proteins triggers syncytia formation. Type-I interferon (IFN)-induced transmembrane proteins (IFITMs), a family of restriction factors that block the entry of many viruses, inhibit S-mediated fusion, with IFITM1 being more active than IFITM2 and IFITM3. On the contrary, the TMPRSS2 serine protease, which is known to enhance infectivity of cell-free virions, processes both S and ACE2 and increases syncytia formation by accelerating the fusion process. TMPRSS2 thwarts the antiviral effect of IFITMs. Our results show that the pathological effects of SARS-CoV-2 are modulated by cellular proteins that either inhibit or facilitate syncytia formation.

**One Sentence Summary:** Syncytia produced by SARS-CoV-2 infected cells and regulation of their
formation by IFITMs and TMPRSS2.

## Introduction

COVID-19 consists of a spectrum of syndromes from mild, flu-like illness to severe pneumonia. Disease severity is linked to lung epithelial destruction, resulting from both immune-mediated damages and viral cytopathic effects. SARS-CoV-2 infection of respiratory epithelial cells likely activates monocytes, macrophages, and dendritic cells, resulting in secretion of proinflammatory cytokines ^1,2^. Excessive systemic cytokine production may lead to thrombosis, hypotension, acute respiratory distress syndrome (ARDS) and fatal multi-organ failure. The innate type-I and type-III interferon (IFN) response, which normally controls viral replication is also reduced in severe cases ^3,4^. However, prolonged IFN-production aggravates disease by impairing lung epithelial regeneration ^5,6^. In the lung, SARS-CoV-2 infects ciliated cells in the airway, alveolar type 2 pneumocytes and epithelial progenitors among others ^7-9^. SARS-CoV-2 and other coronaviruses are cytopathic ^10-14^. The death of infected cells is also a trigger of immune activation.

SARS-CoV-2 entry into cell is initiated by interactions between the spike glycoprotein (S) and its receptor, angiotensin-converting enzyme 2 (ACE2), followed by S cleavage and priming by the cellular protease TMPRSS2 or other surface and endosomal proteases ^15-17^. The structure of S in complex with ACE2 has been elucidated ^18-20^. S consists of three S1-S2 dimers, displaying conformational changes upon virus entry leading to fusion. Besides fusion mediated by virions, S proteins present at the plasma membrane can trigger receptor-dependent syncytia formation. These syncytia have been observed in cell cultures and in tissues from individuals infected with SARS-CoV-1, MERS-CoV or SARS-CoV-2 ^21-28^, but they were not precisely characterized. It has been proposed that they may originate from direct infection of target cells or from the indirect immune-mediated fusion of myeloid cells. Fused pneumocytes expressing SARS-CoV-2 RNA and S proteins were observed in post-mortem lung tissues of 20 out of 41 COVID-19 patients, indicating that productive infection leads to syncytia formation, at least in critical cases ^28^.

SARS-CoV2 replication is in part controlled by the innate host response, through mechanisms that are currently being unveiled. Interferon-stimulated genes (ISGs) inhibit discrete steps of the viral life cycle. At the entry level, the interferon (IFN)-Induced Transmembrane proteins (IFITM1, IFITM2, or IFITM3) block many viruses by inhibiting virus–cell fusion at hemifusion or pore formation stages ^29^. IFITMs act by modifying the rigidity and/or curvature of the membranes in which they reside ^29-32^. Due to different sorting motifs, IFITM1 is mostly found at the plasma membrane, whereas IFITM2/3 accumulate in the endo-lysosomal compartment after transiting through the surface. IFITMs inhibit SARS-CoV, 229E and MERS-CoV entry, but promote infection by HCoV-OC43, a coronavirus that causes the common cold ^33-38^. IFITMs, as well as other ISGs, including LY6E and Cholesterol 25-hydrolase (CH25H), impair SARS-CoV-2 replication by blocking the fusion of virions ^39-41^. Most of the experiments regarding these ISGs have been performed with single-cycle viral pseudotypes. Little is known about the impact of IFITMs on SARS-CoV-2 induced syncytia formation.

Here, we characterized the mechanisms of SARS-CoV-2-induced cell-cell fusion and examined how syncytia formation is impacted by IFITMs and TMPRSS2.

## Results

We first examined whether SARS-CoV-2 infected cells may form syncytia. To this aim, we derived U2OS cells stably expressing ACE2. We selected this cell line because its flat shape facilitates imaging. We generated U2OS-ACE2 cells carrying a GFP–Split complementation system ^42^, in which two cells separately produce half of the reporter protein, producing GFP only upon fusion (Fig. 1A). These U2OS-ACE2-derived cells, that we termed “S-Fuse” cells, were exposed to various doses of SARS-CoV-2. Video-microscopy analysis showed that syncytia appeared rapidly, starting at 6h post-infection, and grew in size, as bystander cells are incorporated in fused cells (Fig. 1B,C and movie S1). Most of the syncytia end up dying, as assessed by the acquisition of a Propidium Iodide (PI) (Fig. 1B,C and movie S1). The extent of fusion was then quantified by measuring the GFP+ area with an automated confocal microscope. The total area of syncytia within each well correlated with the viral inoculum, indicating that the assay provides a quantitative assessment of viral infection (Fig. 2A). S was expressed by the syncytia, but also by single infected cells that have not yet fused, as assessed by immunofluorescence (Fig. 1D). Flow cytometry on unpermeabilized infected cells further showed that S was present at the surface (Fig. 1E).

**Figure 1.**
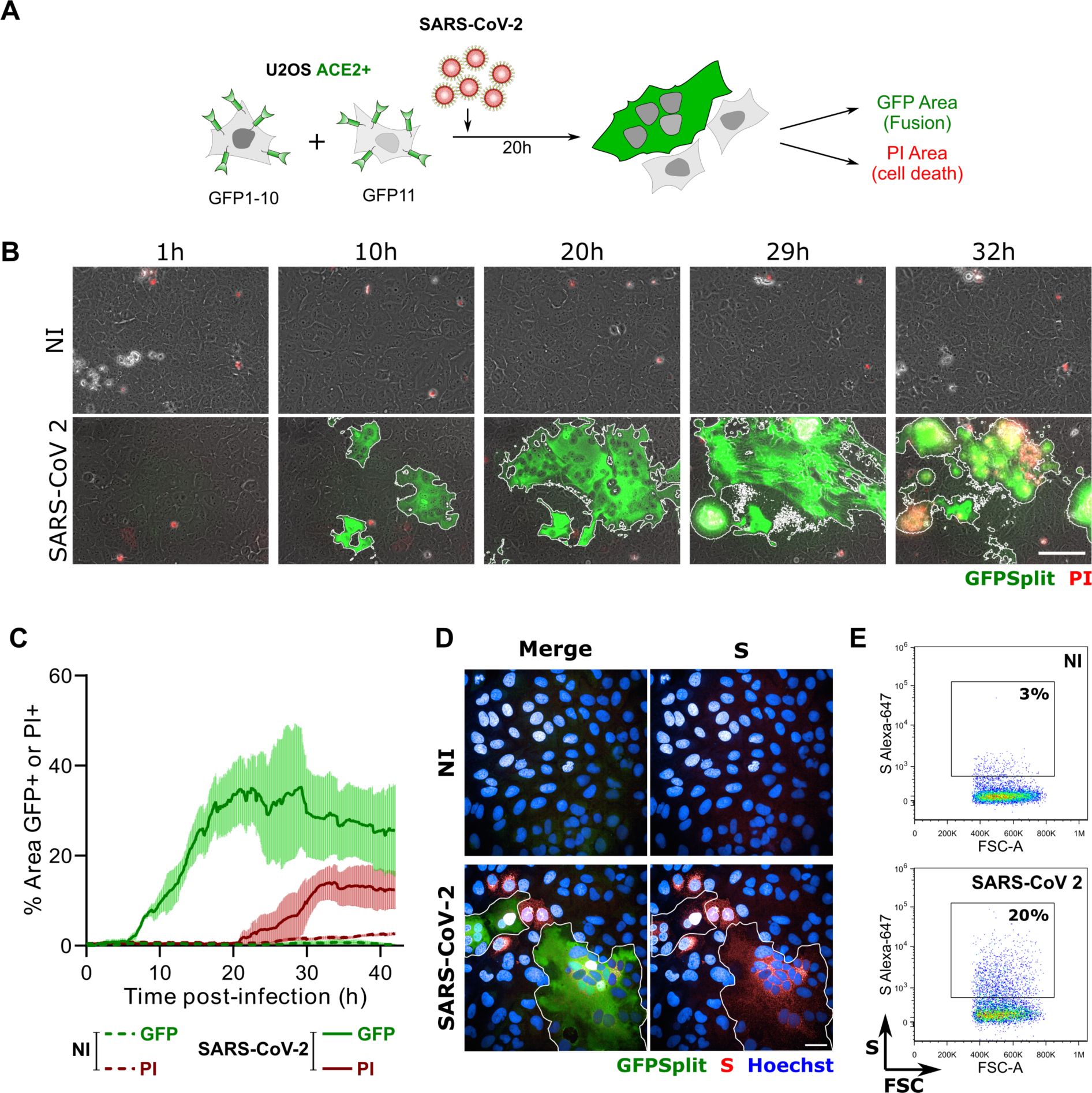
SARS-CoV-2 induced syncytia formation. **A**. GFP-Split U2OS-ACE2 were co-cultured at a 1:1 ratio and infected with SARS-CoV-2. Syncytia formation and cell death was monitored by video microscopy or at endpoint using confocal microscopy and high content imaging. **B**. Still images of GFP (syncytia) and Propidium Iodide (PI) (cell death) at different time-points. Scale bar: 100 µm. **C**. Quantification of U2OS-ACE2 fusion and death by time-lapse microscopy. Results are mean±sd from 3 fields per condition. **D**. S staining of infected U2OS-ACE2 cells analyzed by immunofluorescence. The Hoechst dye stains the nuclei. Scale bar: 40 µm. **E**. Surface S staining of infected U2OS-ACE2 cells analyzed by flow cytometry. Results are representative of at least three independent experiments.

**Figure 2.**
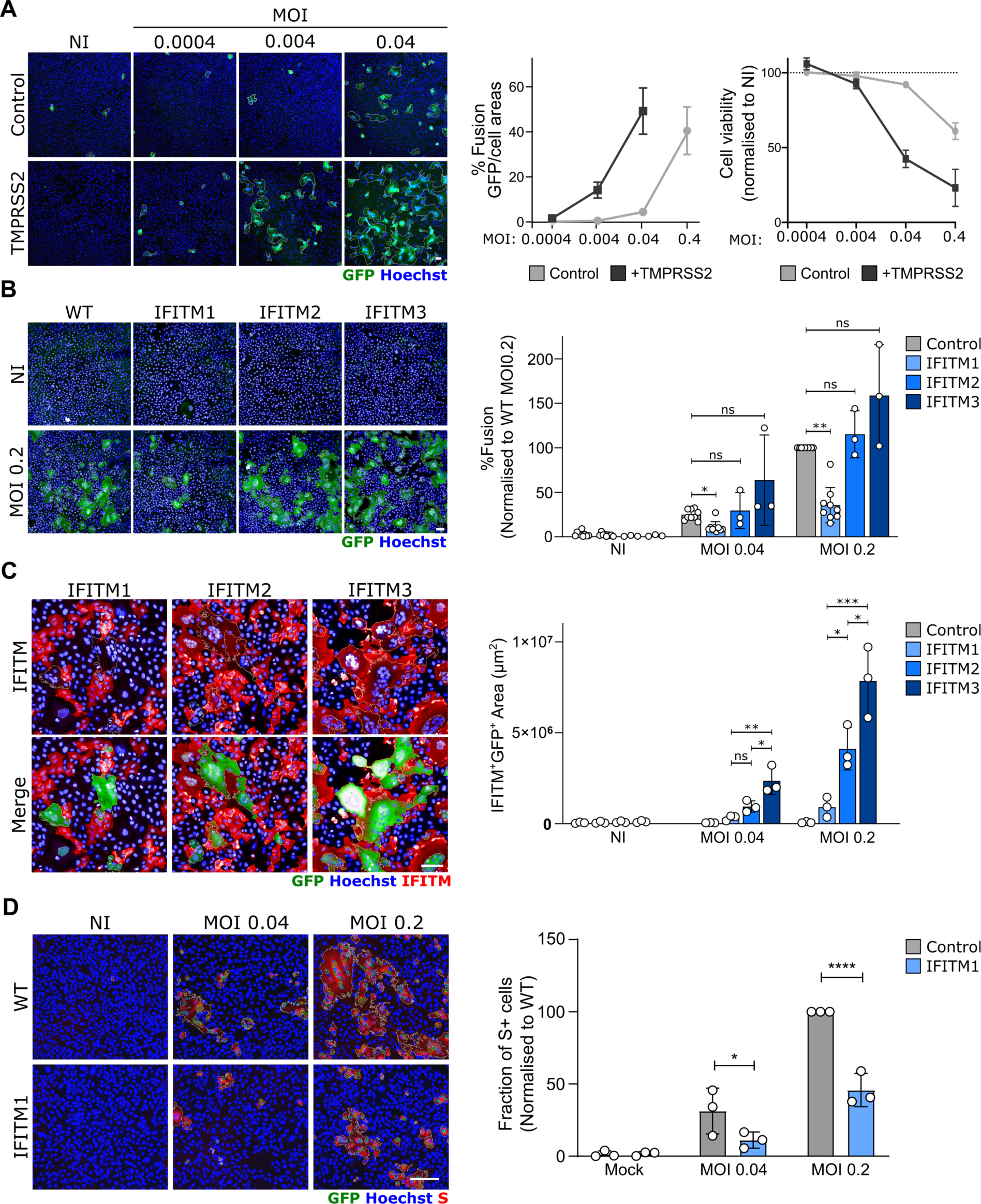
Impact of TMPRSS2 and IFITMs on syncytia formation by U2OS-ACE2 infected cells. Cells were infected at the indicated multiplicity of infection (MOI) and analyzed after 20h. **A**. TMPRSS2 increases fusion and cell mortality. Right panel: Areas of GFP+ cells and PI+ cells **B**. IFITM1, but not IFITM2 and 3, inhibits SARS-CoV-2-induced syncytia formation. Right panels: Area of GFP+ cells **C**. IFITM1+ cells do not fuse with U2OS-ACE2 syncytia. Right panel: Area of IFITM+ GFP+ cells **D**. IFITM1 decreases the number of infected cells. Right panel: Fraction of cells positive for S, normalized to WT cells. Left panels: one representative experiment is shown. Scale bars: 100 µm. Right panels: Data are mean±sd of 3-9 independent experiments. Statistical analysis: B-C: One-Way ANOVA, D: Two-Way ANOVA. ns: non-significant, *p < 0.05, **p < 0.01, ***p < 0.001, ****p < 0.0001.

We then asked whether TMPRSS2 and IFITM1, IFITM2 and IFITM3 impact syncytia formation. We generated S-Fuse cells stably producing each of the four proteins. Their expression was verified by flow cytometry or western blotting (Fig. S1). We then compared their sensitivity to SARS-CoV-2 infection. The presence of TMPRSS2 increased the appearance of fused cells by 5-10 fold (Fig. 2A). On the contrary, IFITM1 significantly inhibited syncytia formation (Fig. 2B and movie S1). Therefore, TMPRSS2 and IFITM1 exert opposite effects on syncytia formation. IFITM2 and 3 were poorly active (Fig. 2B), probably because they mostly accumulate within the endosomal compartment, which limits their ability to alter fusion events occurring at the plasma membrane. Since only 40% of each cell population expressed high levels of IFITM, as assessed by flow cytometry (Fig. S1) and immunofluorescence (Fig. 2C), we asked whether IFITM+ cells were present in the syncytia. A co-staining with anti-IFITM antibodies indicated that syncytia did not incorporate IFITM1+ cells present in the culture, whereas this was not the case for IFITM2 and 3 (Fig. 2C and Fig. S3). Moreover, the inhibitory effect of IFITM1 on syncytia was associated a decreased number of S+ cells (Fig. 2D).

We next assessed whether other cell types form syncytia upon SARS-CoV-2 infection. We generated and infected 293T-ACE2 and A549-ACE2 cells. We also used Vero E6 cells, which are naturally sensitive to infection. The three cell lines actively replicated the virus. Syncytia were readily detected in 293T-ACE2 but not in A549-ACE2 or Vero E6 cells (not shown). Thus, the ability to form syncytia upon SARS-CoV-2 infection is cell-type dependent.

We further characterized the mechanisms of fusion and its regulation by IFITMs and TMPRSS2. We asked whether S alone was sufficient to trigger fusion by transfecting an expression plasmid in 293-T cells harbouring the GFP-Split system (Fig. 3A). Many large and multinucleated GFP+ cells were detected, when ACE2 was co-expressed (Fig. 3B). S-mediated cell fusion was significantly decreased when cells were co-transfected with flag-tagged IFITM1, 2 or 3 plasmids, compared to a control plasmid (Fig. 3B,C). IFITM1 was slightly more inhibitory than IFITM2 and 3 in this system. The transient expression of TMPRSS2 enhanced fusion by 2.5 fold. Interestingly, when the serine protease was present, the inhibitory effect of IFITMs was no longer visible (Fig. 3B,C). The anti-fusogenic effect of IFITM1 varied with the amount of transfected IFITM1 plasmid, but TMPRSS2 counteracted IFITM1 at all doses (Fig. S4). We next measured the kinetics of S-mediated cell fusion by live video-microscopy, monitoring in real-time the GFP+ area. TMPRSS2 accelerated the appearance of GFP+ cells, indicating that it increases the speed of cell-cell fusion (Movie S2 and Fig. 3D). At 12h post-transfection, the syncytia area was already 4-fold larger than in the control condition (Fig. 3D). A similar kinetic analysis indicated that IFITM1 strongly inhibited fusion, whereas IFITM2 and IFITM3 were less efficient (Fig. 3E). In presence of TMPRSS2, the rapid fusion kinetics were similar with or without IFITMs (Fig. 3E).

**Figure 3:**
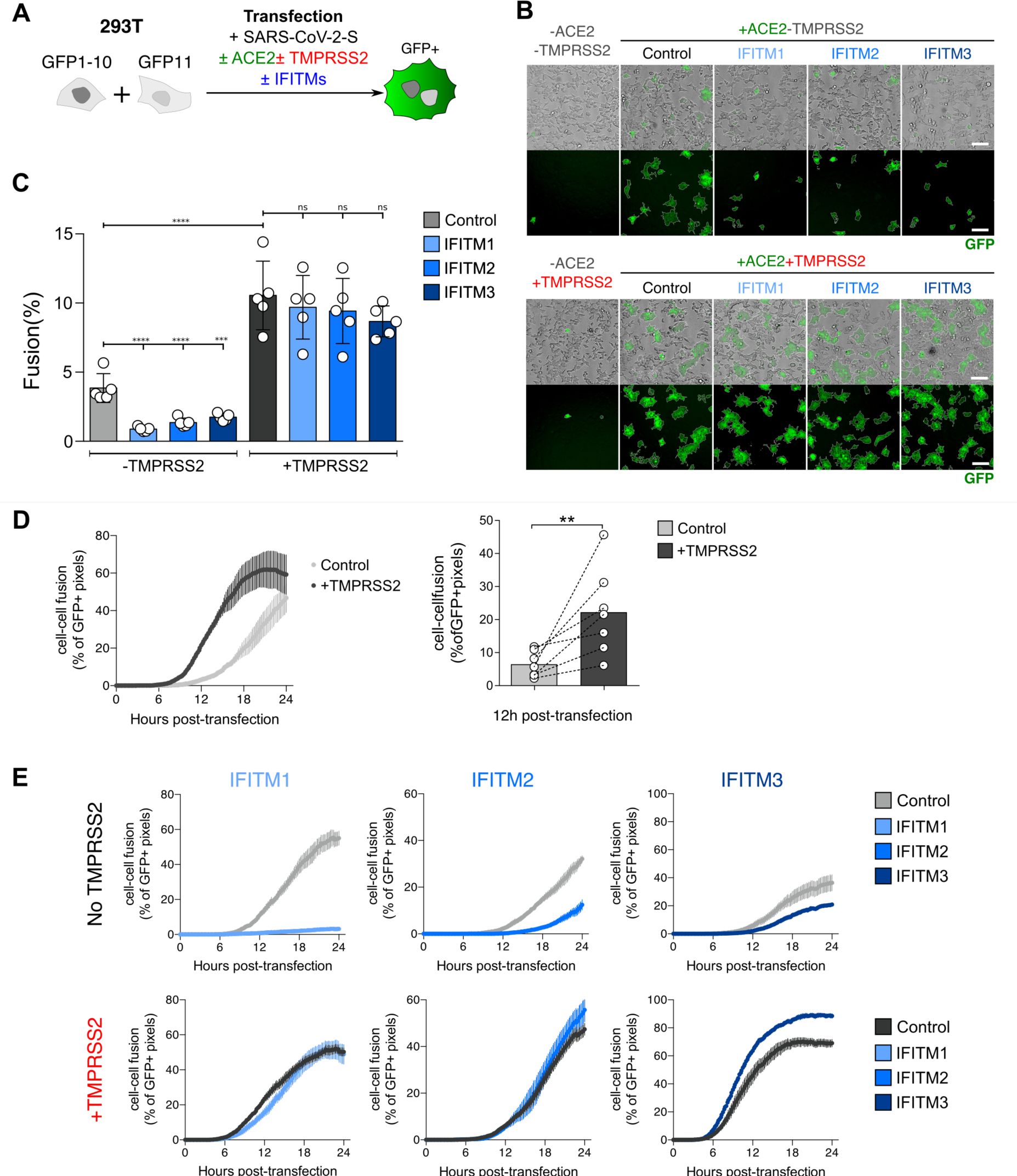
Impact of TMPRSS2 and IFITMs on the kinetics of fusion by S-expressing 293T cells. **A**. 293T-GFP1-10 and -GFP11 cells (1:1 ratio) were co-transfected with S, ACE2, TMPRSS2, IFITM or control plasmids. Cell fusion was quantified by measuring the GFP+ area by high-content imaging after 18h. (B and C) or analyzed over time by video microscopy (D and E). **B**. Representative images of cell-cell fusion. Scale bar: 100 µm. **C**. Quantification of GFP+ areas. Results are mean±sd from 5 independent experiments. **D**. TMPRSS2 accelerates fusion. Cells were monitored by video microscopy and the GFP area was quantified over time. Left panel: one representative experiment. Results are mean±sd from 3 fields per condition. Right panel: Mean±sd from 7 independent experiments (at 12h post transfection). **E**. Impact of TMPRSS2 and IFITMs on the kinetics of fusion by S-expressing 293T cells. One representative out of three independent experiment is shown. Statistical analysis: B, C: One-Way ANOVA, ns: non-significant, ***p < 0.001, ****p < 0.0001. D: Wilcoxon matched-pairs signed rank test

We next studied whether IFITMs and TMPRSS2 impact cell-cell fusion by acting on S-expressing cells (“donor cells”), on ACE2-expressing cells (“acceptor cells”) or on both. To this end, we used a co-culture system of 293T-GFP1-10 donor cells with 293T-GFP11 acceptor cells. IFITMs and TMPRSS2 were transfected into either donor or acceptor cells (Fig. 4A). In the absence of TMPRSS2, IFITMs were poorly efficient when present in donor cells, but inhibited fusion in acceptor cells. As already observed, IFITM1 was more active than IFITM2 and IFITM3. When TMPRSS2 was present in donor cells, IFITMs lost their weak effect they displayed when they were also in donor cells, but retained their ability to inhibit fusion when expressed in acceptor cells (Fig. 4A). Finally, when TMPRSS2 was expressed in acceptor cells, inhibition of fusion by IFITM was abolished regardless of their side of expression (Fig. 4A). Therefore, IFITMs and TMPRSS2 modulate the efficiency of fusion when present in the same cell as ACE2, rather than in the S-expressing cell.

**Figure 4:**
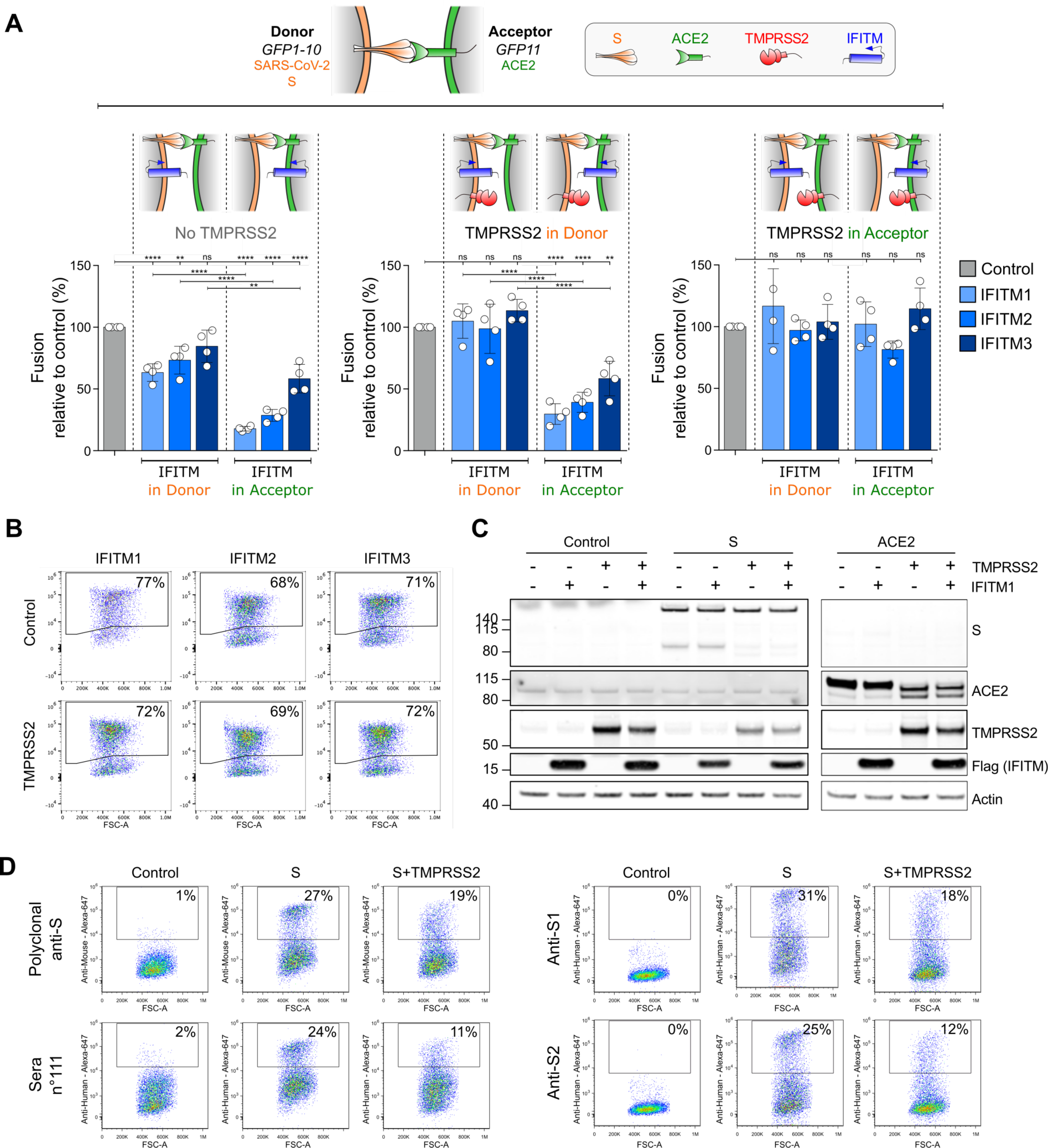
Effect of TMPRSS2 and IFITM on S- or ACE2-expressing cells. **A**. S-expressing cells (Donor cells), ACE2-expressing cells (Acceptor cells) were co-transfected with TMPRSS2, IFITM or control plasmids. Cell fusion was quantified by measuring the GFP+ area after 18h. The indicated combinations were tested. The impact of IFITMs was measured in absence of TMPRSS2 (left panel), in presence of TMPRSS2 in donor (middle panel) or acceptor cells (right panel). Data are mean±sd of 4 independent experiments. Statistical analysis: One-Way ANOVA, ns: non-significant, **p < 0.01, ****p < 0.0001. **B-D**. Impact of TMPRSS2 on IFITMs, ACE2 and S levels. **B**. TMPRSS2 does not decrease IFITM amounts measured by flow cytometry. 293T cells were transfected with the indicated IFITM plasmids, with or without TMPRSS2, and analyzed 18h later. **C**. Impact of TMPRSS2 on S and ACE2, measured by Western Blot. 293T cells were transfected with or without IFITM1, TMPRSS2, S or ACE2 plasmids, and analyzed 18h later. **D**. TMPRSS2 decreases S surface levels, measured by flow cytometry. 293T cells were transfected with S plasmid, with or without TMPRSS2, and analyzed 18h later. Murine polyclonal anti-S antibodies, one serum from a convalescent COVID-19 patient (sera n°111), and two monoclonal antibodies (anti-S1 and anti-S2) were tested. Data are representative of three independent experiments.

Our observation that TMPRSS2 counteracts the inhibitory effect of IFITMs prompted us to examine whether it degrades these antiviral proteins. Flow cytometry and Western blot showed that this is not the case, since similar levels of IFITM1, IFITM2 or IFITM3 were detected in the presence or absence of the serine protease (Fig. 4B). TMPRSS2 decreased S surface levels measured by flow cytometry with antibodies targeting the S1 and S2 domains (Fig. 4D), as well as its processing, visualized by the disappearance of a 90 kDa product by Western Blot (Fig. 4C). In the absence of S, TMPRSS2 also triggered ACE2 processing, a phenomenon which was not inhibited by IFITM1 (Fig. 4C). ACE2 cleavage by TMPRSS2 and other cellular proteases was previously reported and proposed to generate a receptor enhancing viral uptake ^43-45^. Altogether, our results indicate that TMPRSS2 processes both ACE2 and S, but does not degrade IFITMs.

## Discussion

We report here that some SARS-CoV-2 infected cells form large syncytia in culture. This phenomenon occurs in U2OS-ACE2 and 293T-ACE2 cells, but not in two other lines, A549-ACE2 and Vero E6 cells. Syncytia formation is thus a cell type-dependent process and likely relies on parameters such as the amount of cellular or viral proteins involved in fusion, or intrinsic biophysical properties of the membranes. S is expressed at the surface of infected cells and is sufficient to generate fusion with neighbouring cells. IFITMs inhibit syncytia formation, with IFITM1 being the most efficient. IFITM2 and IFITM3 did not inhibit fusion of infected U2OS-ACE2 cells but were partly active in 293-T cells expressing only S. This may be due to the intracellular location of each IFITM, or the high levels of IFITMs generated by transient transfection. Future experiments with IFITM mutants lacking sorting signals or motifs involved in palmitoylation and other post translational modifications will help understanding how these proteins impact SARS-CoV-2 fusion.

We further show that TMPRSS2 accelerates SARS-CoV-2-mediated cell-cell fusion. TMPRSS2, through its serine protease activity, is known to cleave both S and ACE2. TMPRSS2 enhances infectivity and fusogenic activity of different coronaviruses, including HCoV-229E, MERS and SARS-CoV-1 ^22,38,44,46- 48,49^. With SARS-CoV-2, a first cleavage of S by furin at the S1/S2 site is required for subsequent cleavage by TMPRSS2 at S2’ site ^25^. ACE2 cleavage by TMPRSS2 at a polybasic site generates a soluble fragment of the receptor ^43-45^. It has been proposed that this dual action on S and ACE2 facilitates virion uptake by target cells, through mechanisms that are not fully understood. The enhancement of syncytia formation described here with SARS-CoV-2 suggests that TMPRSS2 facilitates a step different from virion uptake. It will be worth determining which structural changes are triggered by TMPRS22 on the viral protein and its receptor, and how these changes may affect the relative affinities of the two proteins and the dynamics of the fusion process.

That TMPRSS2 counteracts the inhibitory activity of IFITMs on SARS-CoV-2-mediated syncytia formation raises interesting questions. This observation is not unprecedented, as two other coronaviruses, HCoV-229E and bat SARS-like WIV1 employ proteolytic pathways to evade IFITM restriction ^38,50^. IFITM proteins modify the rigidity or the lipid content of cellular membranes to prevent fusion. How this biophysical constraint is overcome by TMPRSS2 will require further investigations. For instance, it will be worth assessing the motility of ACE2 and S in membranes, before and during syncytia formation, and the impact of IFITM and TMPRSS2 on these processes. Determining the role of other proteins known to inhibit virion fusion, such as Ly6E and CH25H, and other proteases such as furin will provide a global overview of the mechanisms of cell-cell fusion.

An analysis of 41 post-mortem samples from individuals who died of COVID-19 demonstrated extensive alveolar damage and lung vasculature thrombosis ^28^. In situ hybridization showed the presence of large multinucleated pneumocytes expressing viral RNA and proteins in half of the patients ^28^. Syncytia may thus be considered as a frequent feature of severe COVID-19. It will be worth determining whether the syncytia are also generated in mild cases and whether severe or critical cases are linked to polymorphisms in IFITMs, as already reported for Flu ^29,30^ and in other IFN-related genes. Our results pave the way for the future assessment of the role played by syncytia in viral persistence and dissemination, the destruction of alveolar architecture, and immune or inflammatory responses.

## Acknowledgments

We thank members of the Virus and Immunity Unit for discussions and help, Mauro Giacca for discussion and sharing unpublished results, Nathalie Aulner and the UtechS Photonic BioImaging (UPBI) core facility (Institut Pasteur), a member of the France BioImaging network, for image acquisition and analysis, Nicolas Escriou for the kind gift of anti-SARS-CoV-2 antibodies.

## Funding

OS lab is funded by Institut Pasteur, ANRS, the Vaccine Research Institute (ANR-10-LABX-77), Labex IBEID (ANR-10-LABX-62-IBEID), “TIMTAMDEN” ANR-14-CE14-0029, “CHIKV-Viro-Immuno” ANR-14-CE14-0015-01 and the Gilead HIV cure program, ANR/FRM Flash Covid PROTEO-SARS-CoV-2 and IDISCOVR. Work in UPBI is funded by grant ANR-10-INSB-04-01 and Région Ile-de-France program DIM1-Health. HM lab is funded by the Institut Pasteur, the Milieu Intérieur Program (ANR-10-LABX-69-01), the INSERM, REACTing and EU (RECOVER) grants. SVDW lab is funded by Institut Pasteur, CNRS, Université de Paris, Santé publique France, Labex IBEID (ANR-10-LABX-62-IBEID), REACTing, EU grant Recover. The funders of this study had no role in study design, data collection, analysis and interpretation, or writing of the article.

## Author contribution

JB, JD, MH, BM, DP, MMR, FP, FGB, NC, TB, OS: designed experimental strategy and performed experiments. HM, CP, NE and SVDW provided vital materials and expert advice. JB, OS wrote the manuscript. MMR edited the manuscript. All authors reviewed and approved the final version of the manuscript

## Competing interests

Authors declare no competing interests.

## Data and materials availability

All data associated with this study are available in the main text or the Supplementary Materials. Reagents are available from the authors under material transfer agreements.

## Additional Information

Methods

Fig S1 – S5

Movie S1: Syncytia formation in SARS-CoV-2 infected U20S-ACE2 cells. Left panel: Non infected cells. Middle panel: SARS-CoV-2 infected cells. Right Panel. Inhibition of syncytia formation in SARS-CoV-2 infected U20S-ACE2-IFITM1 cells. Time post infection is indicated

Movie S2: Syncytia formation in S-expressing 293T cells. Left panel: cells transfected with S and ACE2 plasmids. Right Panel. TMPRSS2 accelerates fusion. Cells transfected with TMPRSS2, S and ACE2 plasmids. Time post transfection is indicated

## Methods

### Plasmids

pQCXIP-Empty control plasmid and pQCXIP-IFITM1-N-FLAG, pQCXIP-IFITM2-N-FLAG, pQCXIP-IFITM3-N-FLAG plasmids were described ^42^. pQCXIP-BSR-GFP11 and pQCXIP-GFP1-10 were from Yutaka Hata ^51^ (Addgene plasmid #68716; http://n2t.net/addgene:68716; RRID:Addgene_68716 and Addgene plasmid #68715; http://n2t.net/addgene:68715; RRID:Addgene_68715). pcDNA3.1-hACE2 was from Hyeryun Choe ^52^ (Addgene plasmid # 1786; http://n2t.net/addgene:1786; RRID:Addgene_1786). pCSDest-TMPRSS2 was from RogerReeves ^53^ (Addgene plasmid # 53887; http://n2t.net/addgene:53887; RRID:Addgene_53887). pLenti6-H2B-mCherry was from Torsten Wittmann ^54^ (Addgene plasmid # 89766; http://n2t.net/addgene:89766; RRID:Addgene_89766). pLenti6-attB-hACE2-BSD was generated by cloning hACE2 from pcDNA3.1-hACE2 into the pLenti6-H2B-mCherry. Briefly, hACE2 was PCR amplified adding SpeI, XhoI, and attB sites using forward primer (5’-TCC CTC ACT AGT ACA AGT TTG TAC AAA AAA GCA GGC TGC CAC CAT GTC AAG CTC TTC CTG GCT C -3’) and reverse primer (5’-AAA AAA CTC GAG ACC ACT TTG TAC AAG AAA GCT GGG TTT AAG CGG GCG CCA CCT -3’) and cloned into pLenti6-H2B-mCherry using XhoI and SpeI sites. phCMV-SARS-CoV2-S was previously described ^55^. pLV-EF1α-TMPRSS2-IRES-Hygro was generated by gateway cloning of pCSDest-TMPRSS2 into pLV-EF1α-IRES-Hygro-DEST.

### Cells

293T cells, platinum-E retroviral packaging cell line, U2OS cells and derivatives were cultured in DMEM with 10% Fetal Calf Serum (FCS) and 1% Penicilline Streptomycine (PS). VeroE6 cells were maintained in DMEM with 5% FCS and 1% PS. Cell lines transduced with pQCXIP pLenti6 or pLV-IRES-Hygro derived vectors were grown with 1 μg/ml puromycin, 10 μg/ml blasticidin or **100** μg/ml hygromycin B, respectively (InvivoGen).

### Lentiviral and Retroviral vectors

For lentiviral production, 293T cells were co-transfected with pLenti6 or pLV derived vectors, packaging plasmid R8-2 and VSV-G plasmid as previously described ^42^. For murine retroviral vector production, the platinum-E (Cell Biolabs) retroviral packaging cell line was transfected with pQCXIP-derived plasmids. Vector containing supernatants were harvested at 36h, 48h and 72h and ultracentrifuged 1h at 4°C at 22000 g.

### Generation of stable cell lines

For lentiviral or retroviral transduction, 2×10^4^ cells were resuspended in 150 µl of medium containing 5-25 μl of ultracentrifuged lentiviral or retroviral vectors. Cells were agitated 30 seconds every 5 minutes for 2h30 at 37°C in a Thermomixer. For cell lines co-expressing IFITMs and other proteins, vectors expressing the other proteins were transduced before IFITM vectors to avoid restriction of vector transduction by IFITMs. All cell lines were routinely tested for mycoplasma and found negative.

### GFP-Split fusion assay

For fusion assays with S-expressing cells, 293T-GFP1-10 and 293T-GFP11 cells (6×10^4^ cells/well cells mixed at a 1:1 ratio) in 96 well plates (μClear, #655090) were transfected in suspension using Lipofectamine2000 (Thermo) with 100 ng of DNA. 10 ng of phCMV-SARS-CoV2-S, 25 ng of pCDNA3.1-hACE2, 25 ng of pCSDest-TMPRSS2 and 40 ng of pQCXIP-Empty or pQCXIP-IFITM-N-FLAG were used and adjusted to 100 ng DNA with pQCXIP-Empty. 18 h post transfection, 21 images, covering 90% of the well surface, were acquired per well on an Opera Phenix High-Content Screening System (PerkinElmer) and the cell confluence area and GFP area was quantified on Harmony High-Content Imaging and Analysis Software. For “donor/acceptor” experiments, 3×10^4^ 293T GFP1-10 and GFP11 expressing cells were separately transfected with 50 ng of DNA in suspension at 37°C shaking 900 rpm for 20 minutes using Lipofectamine2000 (Thermo). For donor cells, 293T-GFP1-10 cells were transfected with 10 ng of phCMV-SARS-CoV2-S, ±10 ng pCSDest-TMPRSS2, ±20 ng pQCXIP-IFITM-N-FLAG and adjusted to 50 ng with pQCXIP-Empty. For acceptor cells, 293T-GFP11 cells were transfected with 10 ng of pCDNA3.1-hACE2 ±10 ng pCSDest-TMPRSS2, ±20 ng pQCXIP-IFITM-N-FLAG and adjusted to 50 ng with pQCXIP-Empty. After transfection, cells were washed and resuspended in DMEM 10% FBS, mixed at a 1:1 ratio in different combinations, plated at 6×10^4^ cells per well in a 96 well plate and imaged 18h post transfection. For live imaging, 2.5×10^5^ GFP1-10 and GFP11-expressing 293T cells, mixed at a 1:1 ratio, were transfected in suspension at 37°C, with a shaking at 900 rpm for 20 minutes using Lipofectamine2000 (Thermo) and 100 ng of DNA (10 ng of phCMV-SARS-CoV2-S, 20 ng of pCDNA3.1-hACE2, 20 ng of pCSDest-TMPRSS2 and 40 ng pQCXIP-IFITM-N-FLAG, adjusted to 100 ng with pQCXIP-Empty). Cells were washed and seeded into a µ-Dish 35 mm Quad dish (ibidi – #80416) with 2.5×10^5^ cells per quadrant. Transmission and fluorescence images were taken at 37°C every 10 min up to 24h using a Nikon Biostation IMQ, with three fields for each condition. The GFP area was quantified on ImageJ.

### Antibodies

Mouse monoclonal antibodies (mAb): IFITM1 (#60074-1-Ig, Proteintech) 1:250 for FACS, and IF; IFITM2/3 (#*66081*-1-Ig, Proteintech) 1:250 for FACS, and IF; ACE2 (AC18F) (#AG-20A-0032-C100 – adipogen). Rabbit polyclonal antibodies: FLAG-Tag DYKDDDDK (D6W5B) (#14793, Cell Signaling) 1:800 for FACS. TMPRSS2 (#HPA035787 - Atlasantibodies) 1 :500. Goat polyclonal antibodies: ACE2 (AF933 – R&D). SARS_Ssd3 702, SARS_Ssd3 369, SARS_Ssd3 293, Ascite Sso14 200705 were kindly gifted by Nicolas Escriou. SARS_Ssd3 702 antibody was used at 0.5 μg/ml for FACS and IF. Human Mab: Anti-SARS-CoV2 monoclonal antibodies 48 and 71 recognize the S1 and S2 domains of the S protein, respectively (Planchais et al, manuscript in preparation). Secondary antibodies coupled with Alexa Fluor 488 or 647 (InvitroGen) were used at 1:500 for FACS and IF.

### Flow cytometry

For intracellular staining, cells were fixed in 4% PFA for 15 to 30 minutes at RT and staining was performed in PBS, 1% BSA, 0.05% sodium azide, 0.05% Saponine. Cells were incubated with primary antibodies, and then with secondary antibodies for 30 minutes at RT. Surface staining was performed before fixation, in PBS 1% BSA. Cells were incubated with primary antibodies, and then with secondary antibodies for 30 minutes at RT. Cells were fixed for 15 minutes in 4% PFA. Cells were acquired on an Attune NxT Flow Cytometer (Thermo Fisher) and data analysed with FlowJo software.

### U2OS viral mediated cell-cell fusion, video microscopy and immunofluorescence

U2OS-ACE2 GFP1-10 or GFP11 stably expressing TMPRSS2, IFITM1, 2, 3 or control cells were mixed (1:1 ratio) and plated at 8×10^3^ cells per well in a 96 well plate (μClear, #655090), 24 h before infection. Cells were then infected with different MOI of SARS-CoV-2 in 150 µl of media. 18h post infection, media was removed, and cells were fixed in 4% PFA for 30 min at room temperature, washed and resuspended in PBS. For fusion and viability quantification, cells were stained for 10 min with Hoechst 33342 (1:5000) before being imaged. For S and IFITM analysis, cells were washed twice PBS and stained with primary antibody for 45 min in PBS, 1% BSA, 0.05% sodium azide, 0.05% Saponine in the wells. Cells were washed twice with PBS, 0.05% Saponine and stained with secondary antibody for 30 min at RT in PBS, 1% BSA, 0.05% sodium azide, 0.05% Saponine and Hoechst 33342 (1:10 000) and washed with PBS. Cells were acquired on an Opera Phenix High Content Screening System (PerkinElmer) and analysed on Harmony High-Content Imaging and Analysis Software. For live imaging, U2OS-ACE2 GFP1-10 and GFP11 were mixed at a 1:1 ratio and plated at 4×10^4^ cells per quadrant in a µ-Dish 35 mm Quad dish (ibidi – #80416). Cells were infected the next day with SARS-CoV-2 (MOI 0.2) in media containing Propidium Iodide. Transmission and fluorescence images were taken at 37°C every 10 min, up to 48 h, using a Nikon Biostation IMQ, with three fields for each condition.

### Western Blot

Cells were lysed in TXNE buffer (1% Triton X-100, 50 mM Tris-HCl (pH 7.4), 150 mM NaCl, 5 mM EDTA, protease inhibitors) for 30 min on ice. Equal amounts (20-50 lll g) of cell lysates were analyzed by Western Blot. The following antibodies were diluted in WB-buffer (PBS, 1% BSA, 0.05% Tween, 0.01% Na Azide): goat anti-human ACE2 (R&D cat#AF933, 1:2000), mouse anti-human ACE2 (Adipogen AC18F cat #AG-20A0032-C100, 1:1000), rabbit anti-human TMPRSS2 (Atlas antibodies cat# HPA035787, 1:1000), mouse anti-Flag tag (Sigma cat# F1804, 1:1000), rabbit anti-human actin (Sigma cat#A2066, 1:2000), mouse ascite anti-SARS S, (1:1000) {Siu, 2008 #4573}. Specie-specific secondary DyLight-coupled antibodies were used (diluted 1:10000 in WB-buffer) and proteins revealed using a Licor Imager. Images were quantified and processed using ImageStudioLite software.

### Virus

The strain BetaCoV/France/IDF0372/2020 was supplied by the National Reference Centre for Respiratory Viruses hosted by Institut Pasteur (Paris, France) and headed by Pr. S. van der Werf. The human sample from which the strain was isolated has been provided by Dr. X. Lescure and Pr. Y. Yazdanpanah from the Bichat Hospital, Paris, France. The viral strain was supplied through the European Virus Archive goes Global (Evag) platform, a project that has received funding from the European Union’s Horizon 2020 research and innovation program under grant agreement n° 653316. Titration of viral stocks was performed on Vero E6, with a limiting dilution technique allowing a calculation of DCP50.

### Statistical analysis

Flow cytometry data were analyzed with FlowJo v10 software (TriStar). Calculations were performed using Excel 365 (Microsoft). Figures were drawn on Prism 8 (GraphPad Software). Statistical analysis was conducted using GraphPad Prism 8. Statistical significance between different groups was calculated using the tests indicated in each figure legend.

## Supplementary Figures

**Fig. S1.**
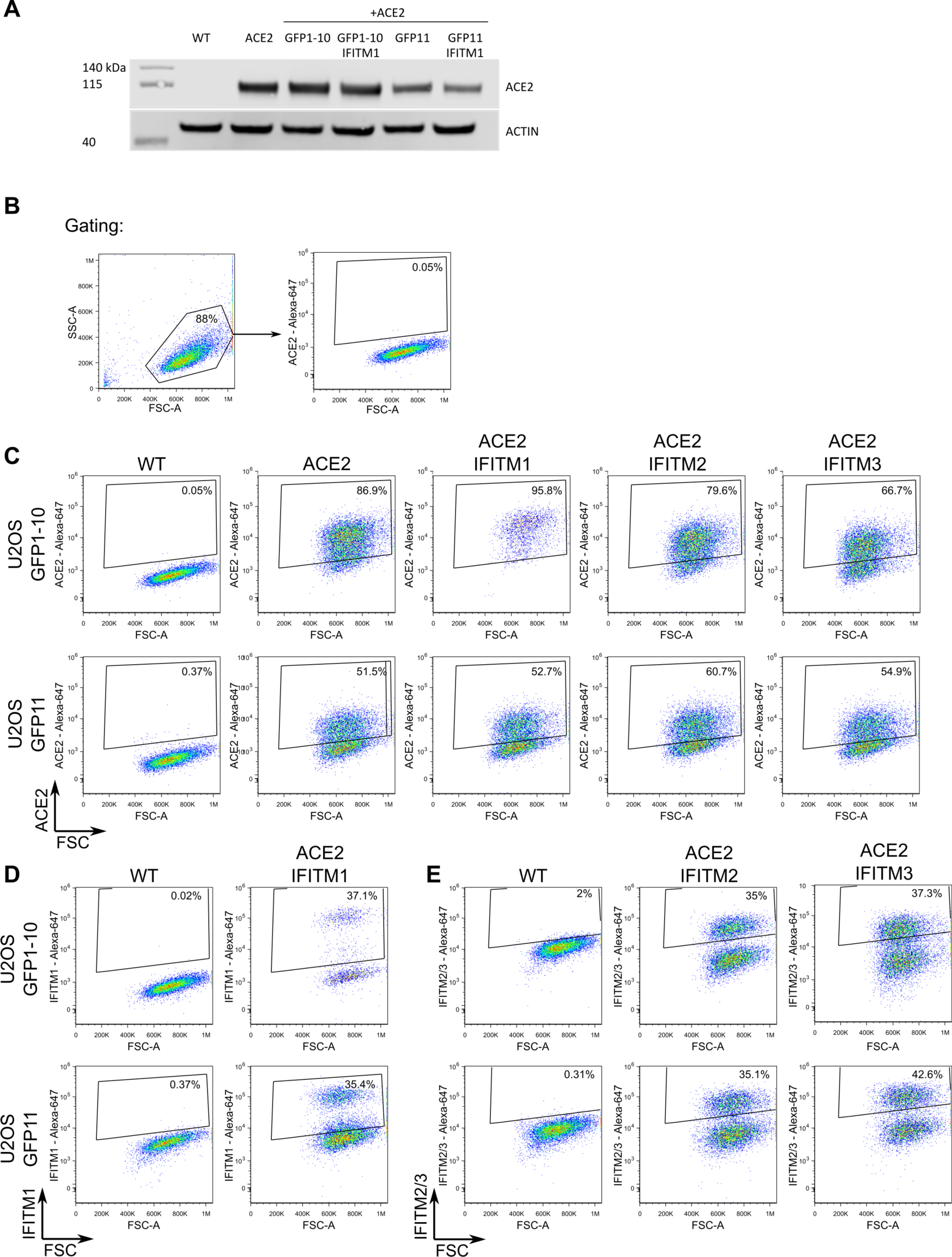
ACE2 and IFITM expression in U2OS-ACE2 cell derivatives. **A**. ACE2 expression assessed by western blot in U2OS-ACE2 GFP-split cells expressing or not IFITM1. Cell lysates were analysed by for ACE2 and actin as a loading control. **B-E**. Analysis of ACE2 and IFITMs levels by flow cytometry **B**. Gating strategy. Cells were gated by size and granularity. Positive and negative gates were then set on control U2OS cells lacking the protein of interest. **C**. ACE2 levels on parental U2OS (WT) and various IFITM derivatives. **D**. IFITM1 levels on parental U2OS (WT) and U2OS-ACE2 IFITM1 cells. **E**. IFITM2 and IFITM3 levels on parental U2OS (WT) and U2OS-ACE2 IFITM2 and U2OS-ACE2 IFITM3 cells.

**Fig. S2.**
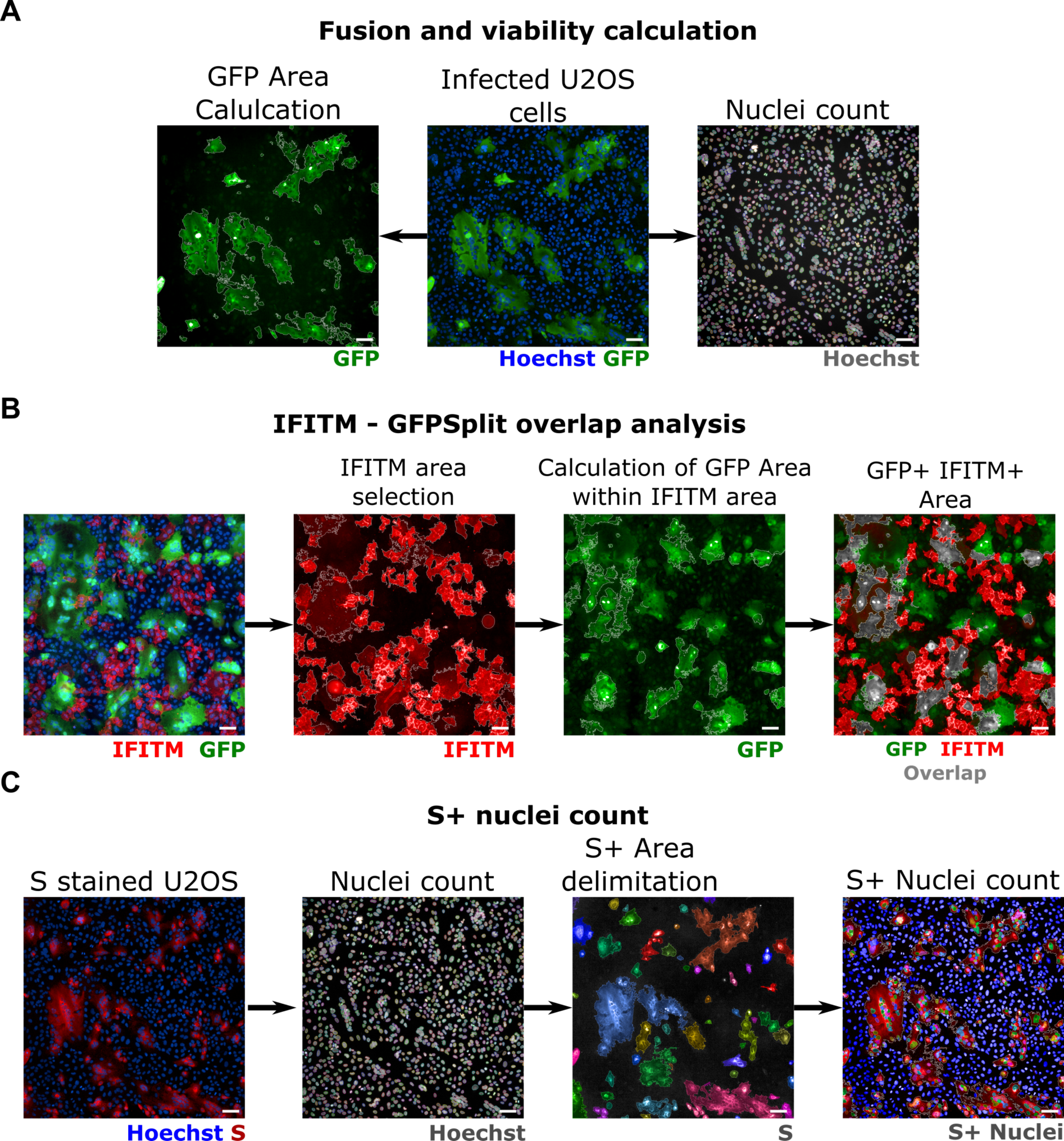
Image quantification methodology. **A**. Quantification of fusion and viability. To measure the extent of cell-cell fusion, the GFP area was automatically delimited, measured and then divided by the total cell area. For cell viability, nuclei were automatically counted, and the total number of nuclei per well was normalized to that of non-infected control cells. **B**. Quantification of syncytia expressing IFITMs. For IFITM-GFP overlap quantification, the IFITM+ area was first selected on the 647 nm channel (image 2) and the GFP positive area was quantified within the IFITM+ area (image 3). Image 4 shows the overlap area in grey for simpler visualisation. **C**. Quantification of infected cells expressing the S protein. The cells were stained with anti-S antibodies and the nuclei were detected with Hoechst. The S+ area was delimited (each selected object is pseudo-coloured). Nuclei present within the S+ area were scored and divided by the total number of nuclei, to calculate the number of infected cells per well. Scale bars: 100 µm.

**Fig. S3.**
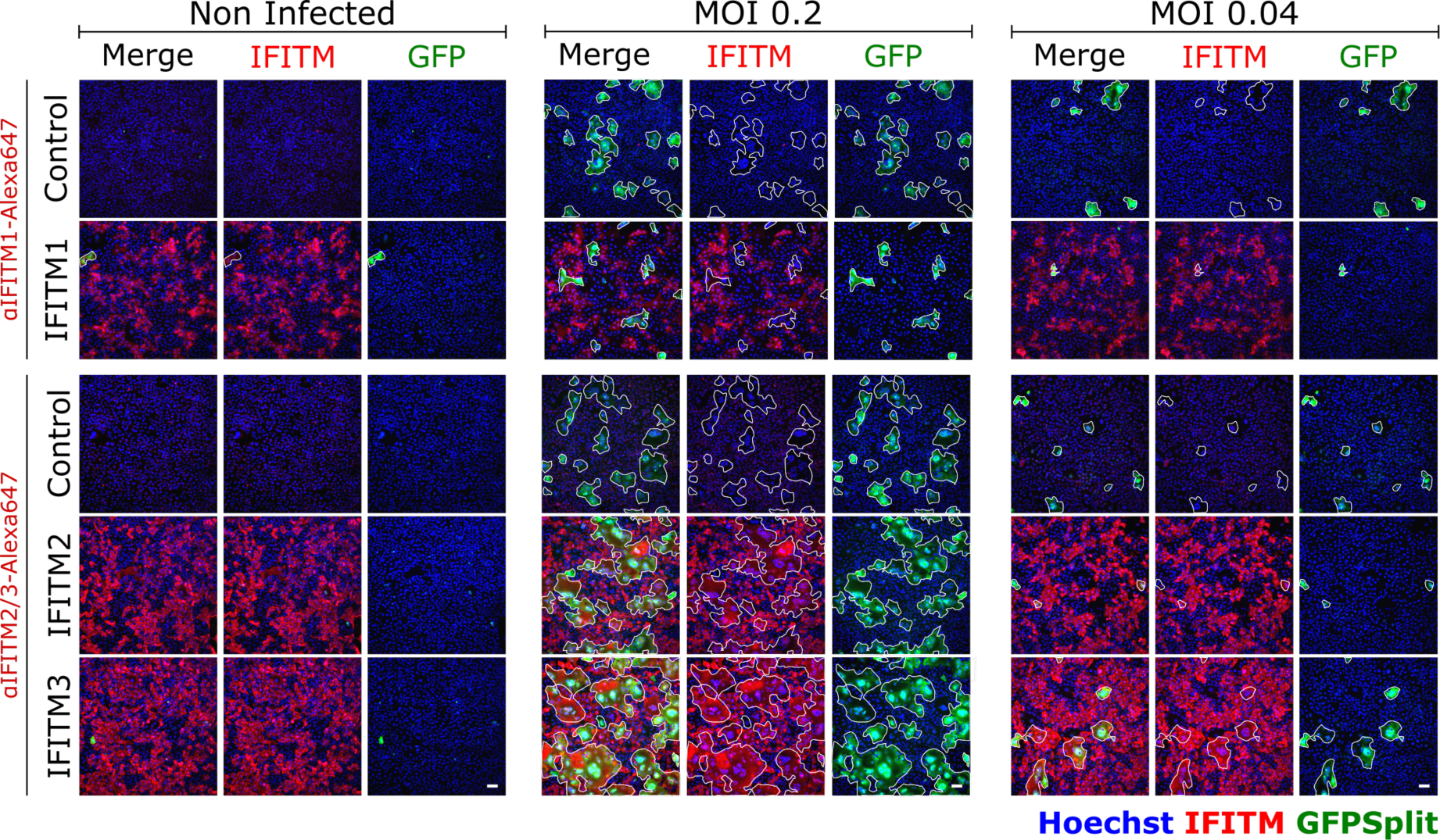
Expression of IFITM1, IFITM2 and IFITM3 in SARS-CoV-2 infected syncytia. in SARS-CoV-2 induced syncytia in U2OS cells are IFITM1 dim or negative, but IFITM2/3 positive. Left: Non infected U2OS-ACE2 cells. Middle and right: Cells infected with SARS-CoV-2 at the indicated MOI. The GFP area is delimited in white. With IFITM1, the fused GFP area does not overlap with the red IFITM1 area. For IFITM2 and IFITM3, an overlap with the GFP is visible. Scale bars: 100 µm. Quantifications are presented Fig. 2C

**Fig. S4.**
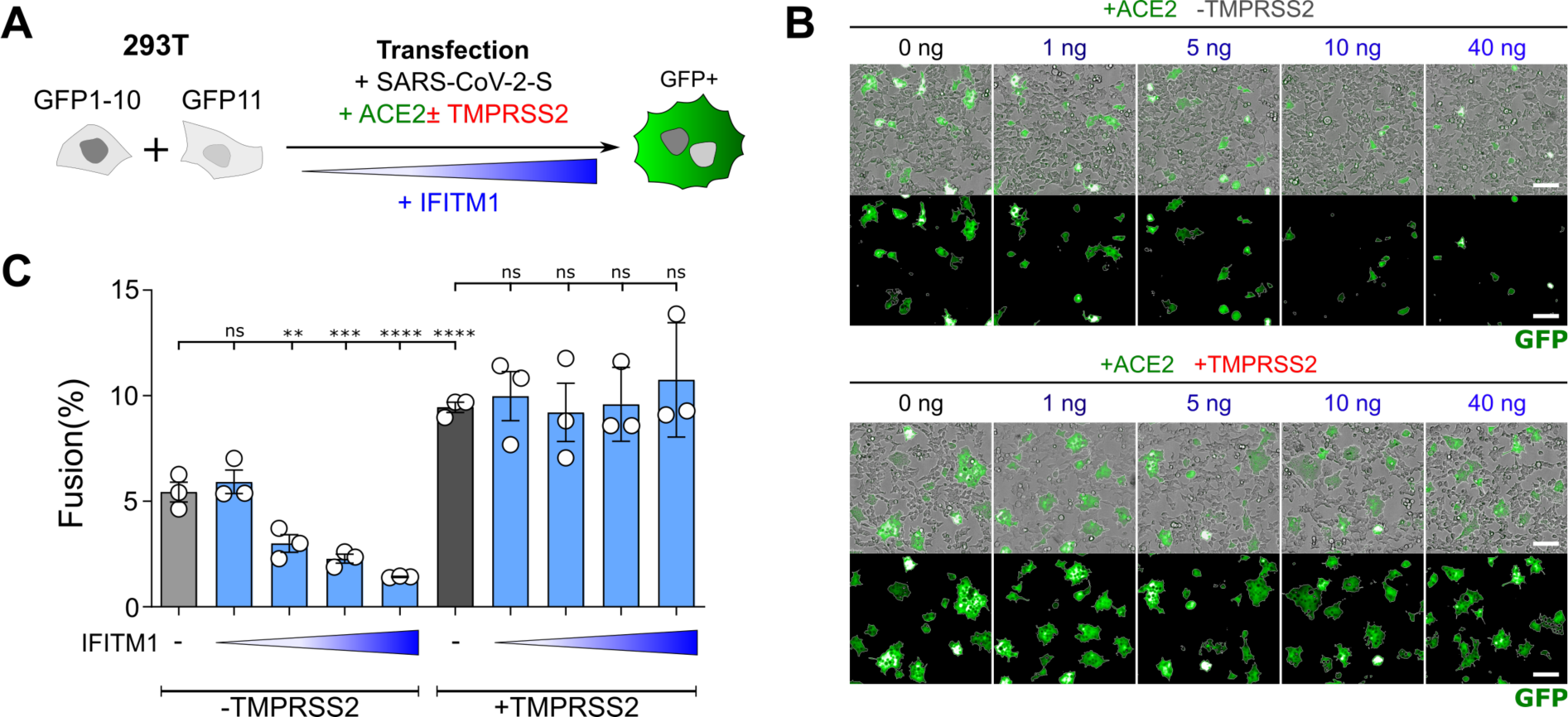
Dose-response analysis of IFITM1 activity. **A**. 293T-GFP1-10 and -GFP11 (1:1 ratio) were co-transfected with S, ACE2, TMPRSS2 and increasing amounts of IFITM1 plasmids. Cell fusion was quantified by measuring the GFP+ area by high-content imaging at 18h post transfection **B**. Representative images of cell-cell fusion, with the indicated amounts of transfected plasmids. Scale bar: 100 µm. **C**. Quantification of GFP+ area. Results are mean±sd from 3 independent experiments. Statistical analysis: One-Way ANOVA, ns: non-significant, **p<0.01, ***p < 0.001, ****p < 0.0001.

**Fig. S5.**
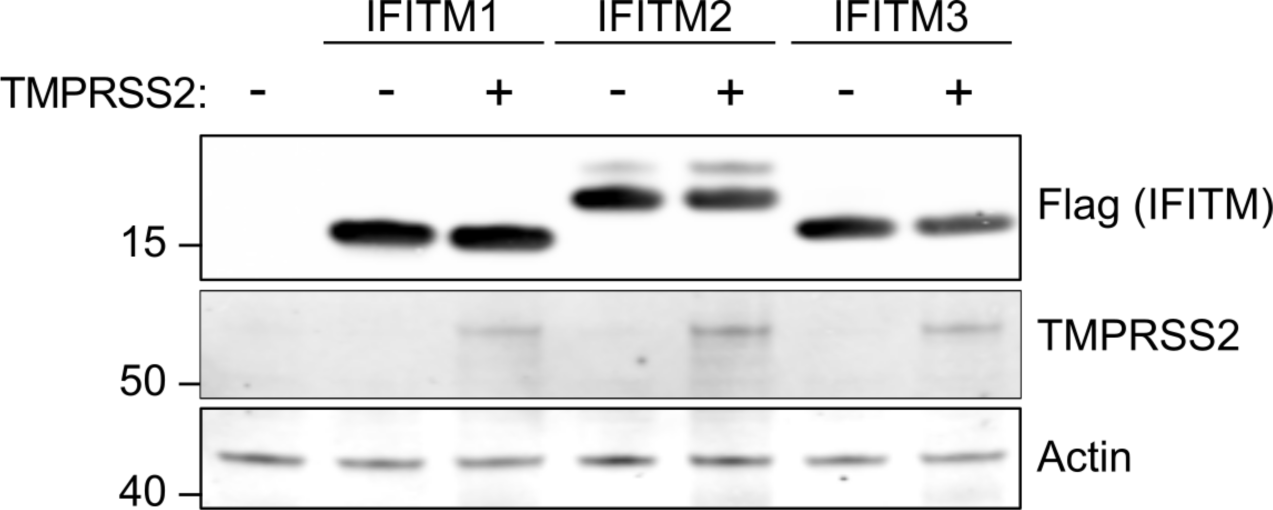
Impact of TMPRSS2 on IFITM1, 2 and 3 levels measured by Western Blot. 293T cells were co-transfected with TMPRSS2 and IFITM1, IFITM2, IFITM3 or control plasmids, and analysed 18h post transfection by western blot. TMPRSS2 does not cleave or reduce the levels of IFITMs.

## Notes

### Competing Interest Statement

The authors have declared no competing interest.

